# *Flavobacterium johnsoniae* Tyrosine Ammonia Lyase (FjTAL) *in-silico* Structure Prediction and Molecular Docking to L-Tyrosine, p-Coumaric Acid (pHCA) and Caffeic Acid

**DOI:** 10.1101/2022.02.09.479702

**Authors:** Seyyed Amirreza Mousavi Majd

## Abstract

Responsible for the conversion of L-tyrosine to p-coumaric acid in *Flavobacterium johnsoniae*, FjTAL has drawn the attention of many biochemical engineers who wish to carry out a sustainable biosynthetic scheme for the production of aromatic compounds. In this study, with the aid of various computational tools, the secondary and tertiary structures of FjTAL have been predicted. The results suggest that FjTAL forms a homo-tetramer when active as a cytosolic enzyme and it is mostly consisted of alpha helices. With the aid of molecular docking, one can hypothesize that FjTAL is likely to bind to L-tyrosine, p-coumaric acid, and caffeic acid with a similar molecular mechanism and thus, p-coumaric acid and caffeic acid may exhibit a negative feedback response toward the enzyme and inhibit its activity competitively. Two distinct binding pockets have been discovered, one of which contains highly conserved residues among several species. The residues which form the prosthetic group 3,5-dihydro-5-methylidene-4H-imidazol-4-one (MIO) also emerge in the evolutionary conserved binding pocket. The other discovered cavity, could either be a second binding site for the ligands or simply an artifact of the molecular docking task.

## 1. Introduction

### 1.1 Background

In the recent methods that have been developed for the sustainable production of phenylpropanoids in recombinant *E. coli*, FjTAL has been proposed as one of the potential enzymes to convert L-tyrosine to p-coumaric acid^1–3^ (Fig.1). Although it has been demonstrated that *Flavobacterium johnsoniae* Tyrosine Ammonia Lyase (FjTAL), has satisfactory affinity for L-tyrosine (K_m_ ~ 6.7 μM), little has been reported on the binding affinity of other structurally-related metabolites to FjTAL^4 5^. It has been hypothesized by Sariaslani^6^ (2007) and Haslinger and Prather (2020) that p-coumaric acid, the product of elimination reaction catalyzed by FjTAL, might inhibit the activity of the enzyme, exerting a kind of negative feedback response. The goal of this study, is to computationally predict the 3D structure of FjTAL, paving the way to help other researchers to reasonably engineer the protein for the future studies. It is worth mentioning that since not even a single experimental data on the structure of the enzyme in Protein Data Bank (PDB) was available, one had to only rely on computational methods in order to determine the structure of the enzyme.

**Fig.1.**
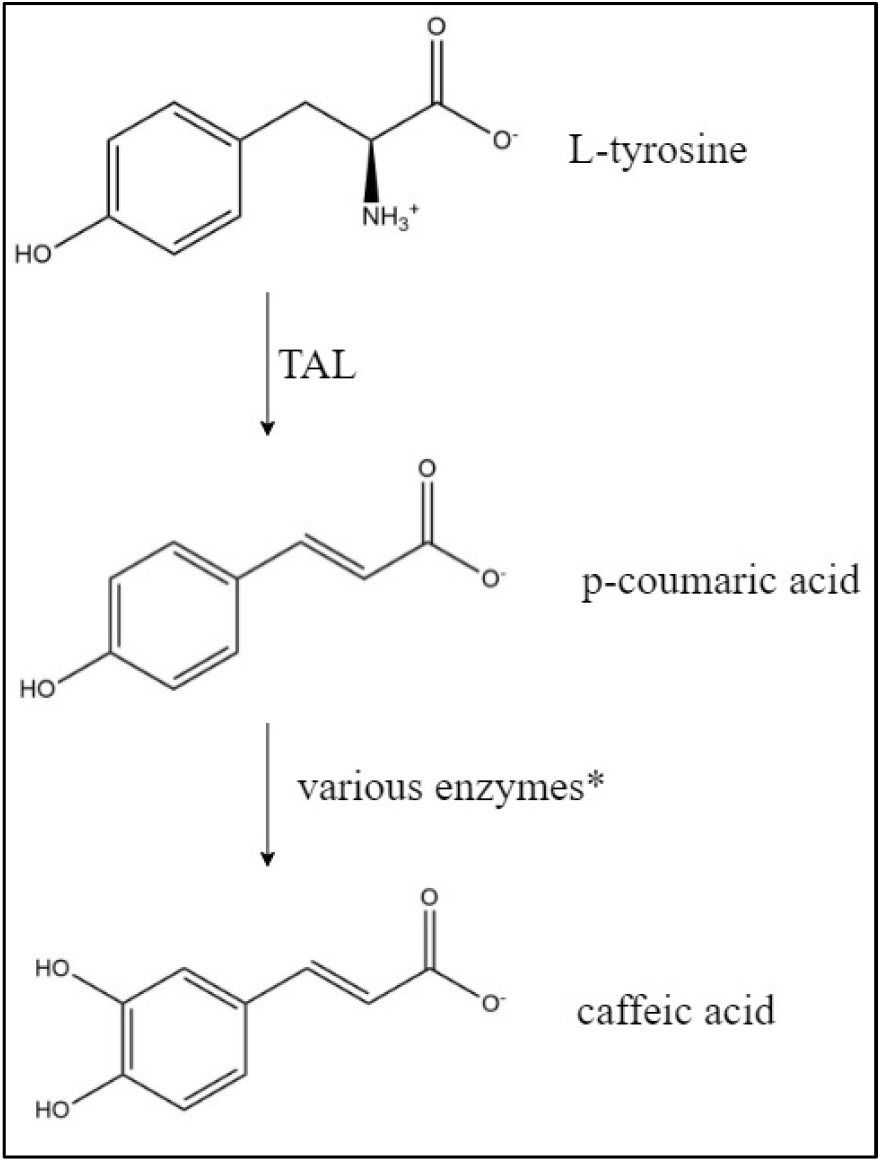
The artificial biosynthetic pathway developed by Rodrigues et al. (2015)^2^ and Haslinger and Prather (2020)^1^ and the structure of related metabolites at pH=7.4 TAL: tyrosine ammonia lyase, e.g., FjTAL, RgTAL, SeSam8, RsTAL, HaTAL *Many different enzymes are utilized for the next step of the artificial pathway including CYP199A2, 4-coumarate 3-hydroxylase, hydroxyphenylacetate 3-hydroxylase

### 1.2 The in-silico Experiment Design

The general strategy devised in this study is depicted in Fig.2. To predict the 3D structure of FjTAL, the sequence of the enzyme had to be retrieved. The codon-optimized DNA sequence of FjTAL was retrieved from the supplementary file of Haslinger and Prather (2020)^1^, and translated it into amino acid alphabet by using the Biostrings package^7^. It was also observed that a record had already existed in NCBI with the exact sequence (SCV44818.1 unnamed protein product).

**Fig.2.**
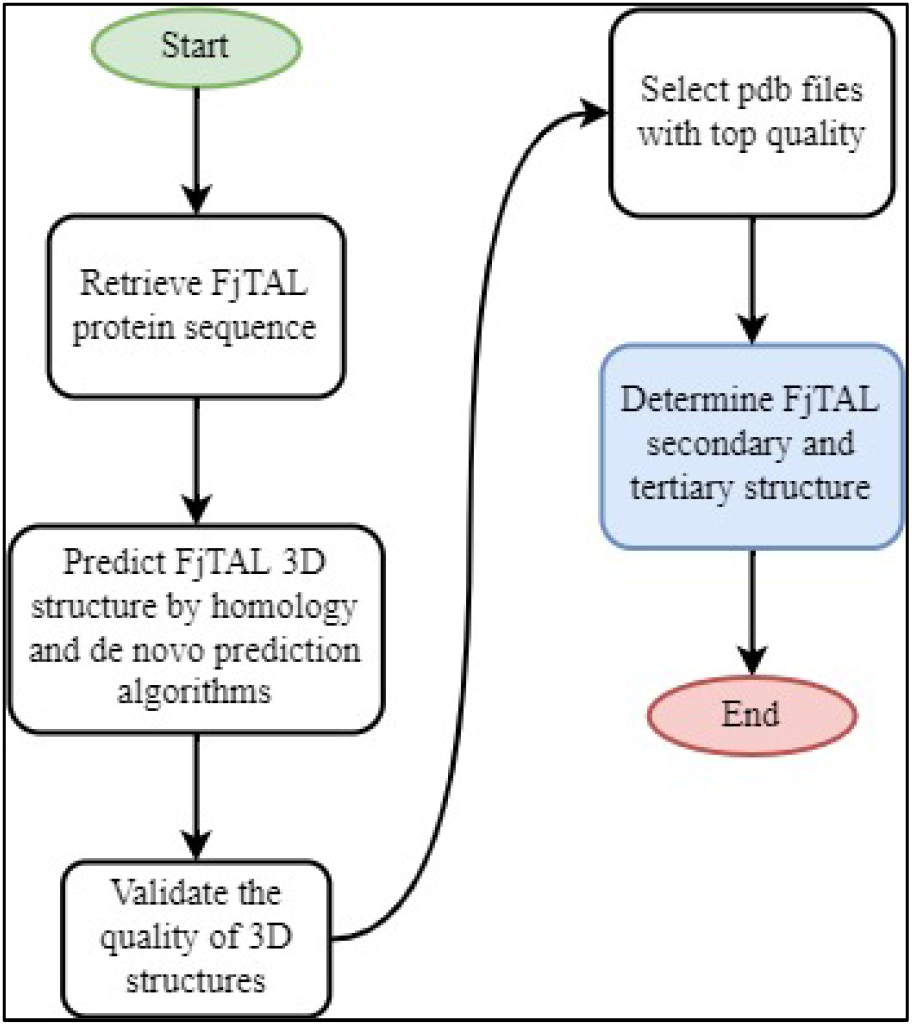
The workflow carried out in this study

With the FjTAL protein sequence available, four webservers were employed in order to predict tertiary structure of FjTAL. Two of these webservers, SWISS-MODEL^8–11^ and Phyre2^12^, implement homology modelling algorithms while the other two, AlphaFold2^13–15^ and RoseTTAFold^16^, predict the structures without the aid of template (ab-initio prediction).

FjTAL protein sequence was fed into each of these four servers. Then, the quality of these structures was validated using the SAVES of UCLA^17–19^ and the ones that did not meet the standards were eliminated. Eventually, the top five pdb files which were found to be superior among all, were stored as .pdb files for the next steps. After that, the secondary structures of the predicted models were extracted from the pdb files and secondary structure prediction methods that were only dependent on the input sequence rather than tertiary structure.

## 2. Materials and Methods

In this study quite a few computational tools were employed to help us fulfil the computational experiment. In brief, the workflow started with employing Biostring package to retrieve the sequence of FjTAL. Then the tertiary structure prediction was done by using Phyre2, SWISS-MODEL as well as ColabFold and Robetta. SAVES of UCLA was helpful in ranking the predicted structures. The molecular docking task (see section 3.3) was executed by CB-Dock and the results were subsequently analyzed by ChimeraX, Mol* and BIOVIA. The complete list of the software and webservers is given in Table1.

**Table1.**
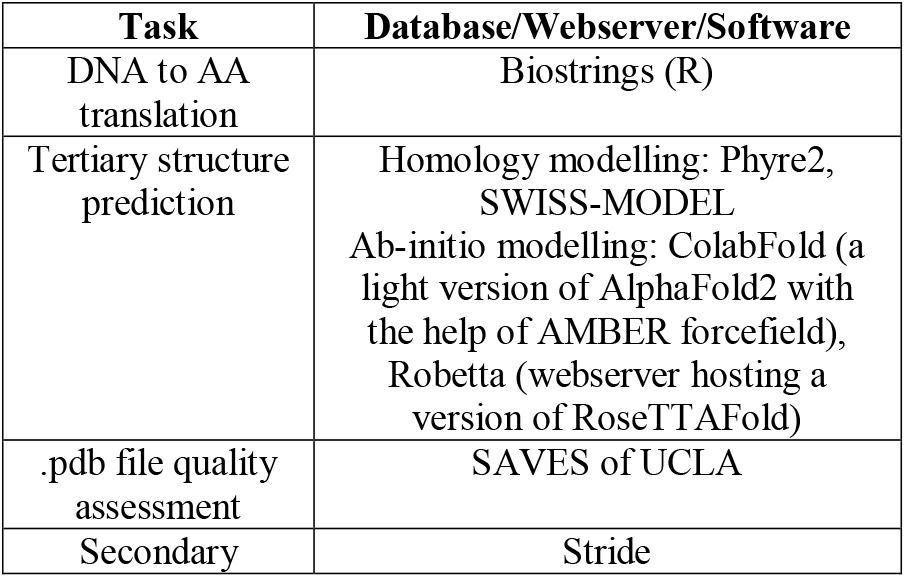
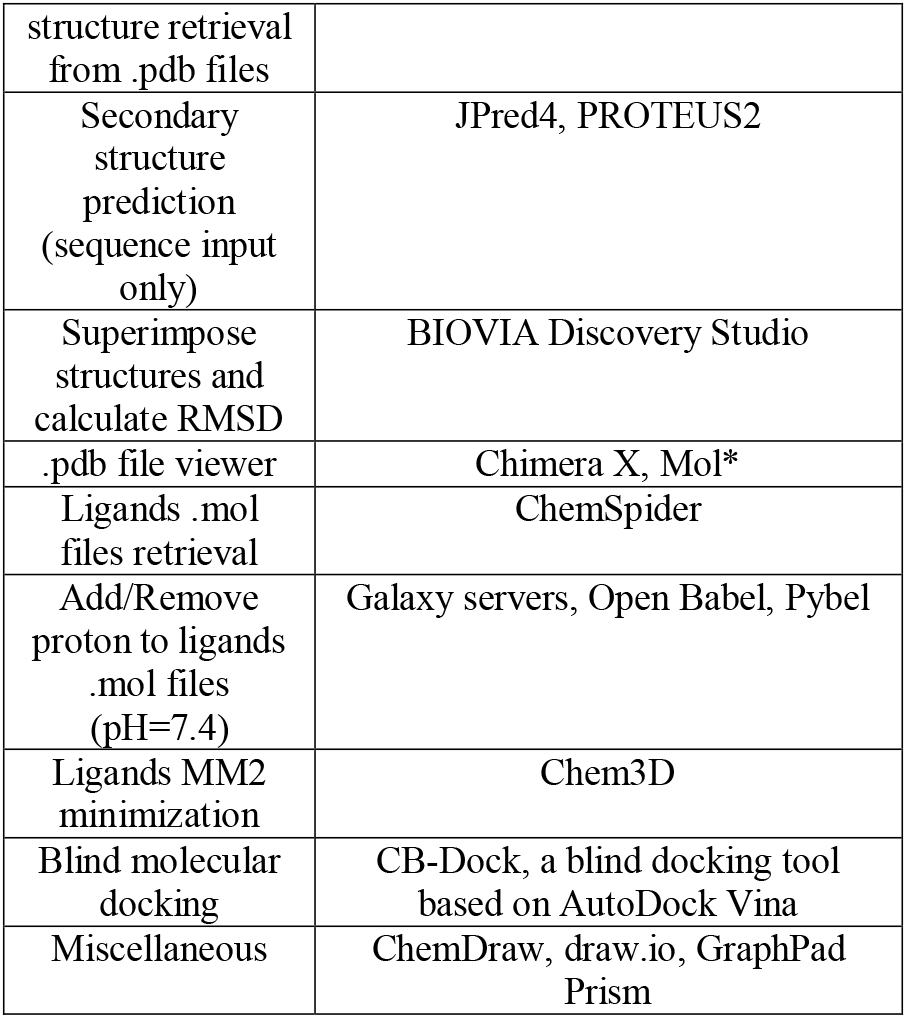
the list of computational tools used throughout this survey.

## 3. Results

### 3.1 FjTAL Structure Prediction and Quality Assesment

As mentioned before, four webservers were employed to predict the structure of FjTAL. Phyre2, SWISS-MODEL, ColabFold and Robetta generate one, two, five and five structures respectively. It is worth noting that ColabFold initially generates five unrelaxed structures and five AMBER-relaxed structures which we used only the latter for the purposes of this study. SWISS-MODEL also predicted that FjTAL is likely to be either a homotetrameric or a homodimeric protein. However, most of the closely-related homologs of FjTAL were proven to be homotetrameric and thus, we have concluded this may also be a valid assumption for FjTAL as well.

All thirteen pdb files, were passed to SAVES webserver to assess the quality of the predicted structures. The ideal structures had to pass Verify3D test and achieve a score > 95 in ERRAT program (top five structures were selected). As shown in Table2, ab-initio algorithms significantly outperformed Phyre2 and SWISS-MODEL. The superiority of AlphaFold2 and RoseTTAFold over classical methods had also been proposed in other studies such as CAMEO^20 21^ as well. The top five structures were selected and stored.

**Table2.**
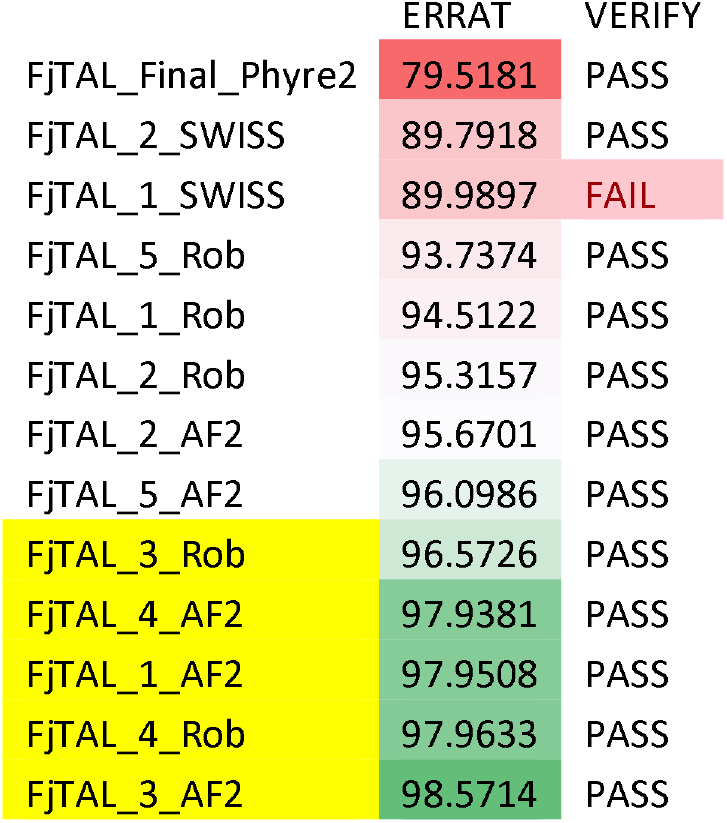
The results of quality assessment using SAVES. FjTAL_Final_Phyre2.pdb is the file generated on Phyre2 server. FjTAL_X_SWISS.pdb files are the output of SWISS-MODEL. FjTAL_X_Rob.pdb files are the tertiary structures predicted by Robetta/RoseTTAFold. FjTAL_X_AF2 files are the output of ColabFold/AlphaFold2.

|  | ERRAT | VERIFY |
| --- | --- | --- |
| FjTAL_Final_Phyre2 | 79.5181 | PASS |
| FjTAL_2_SWISS | 89.7918 | PASS |
| FjTAL_1_SWISS | 89.9897 | FAIL |
| FjTAL_5_Rob | 93.7374 | PASS |
| FjTAL_1_Rob | 94.5122 | PASS |
| FjTAL_2_Rob | 95.3157 | PASS |
| FjTAL_2_AF2 | 95.6701 | PASS |
| FjTAL_5_AF2 | 96.0986 | PASS |
| FjTAL_3_Rob | 96.5726 | PASS |
| FjTAL_4_AF2 | 97.9381 | PASS |
| FjTAL_1_AF2 | 97.9508 | PASS |
| FjTAL_4_Rob | 97.9633 | PASS |
| FjTAL_3_AF2 | 98.5714 | PASS |

### 3.2 FjTAL Secondary and Tertiary Structure

The top models from previous sections (Table2 yellow highlights) were passed to Stride^22^ to extract the secondary structures from pdb files.

We also tried to predict the secondary structures using JPred4^23^ and PROTEUS2^24^ webservers which require the protein sequence as their only input.

In Table3 the prediction results by JPred4, PROTEUS2 and the extraction results by Stride are tabulated.

**Table3.**
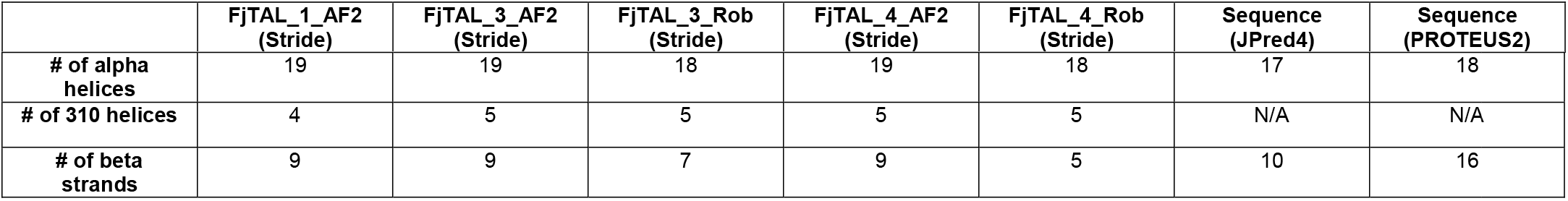
The results of secondary structure prediction and extraction

As shown in Table3, while, to some extents, the results differ from one model to another, one can state that FjTAL is mostly consisted of alpha helices with very few, short length, beta strands.

Note that JPred4 and PROTEUS2 do not generate any data on the number of 310 helices that may exist in the structures and also notice that the number of beta strands differ significantly in each record, mainly because FjTAL seems to have very short (length = < 4 amino acid) beta strands. This has led to emergence of presumably false positive results in the case of beta strands especially in the prediction-based results (See Fig.3 and Fig.4).

**Fig.3.**
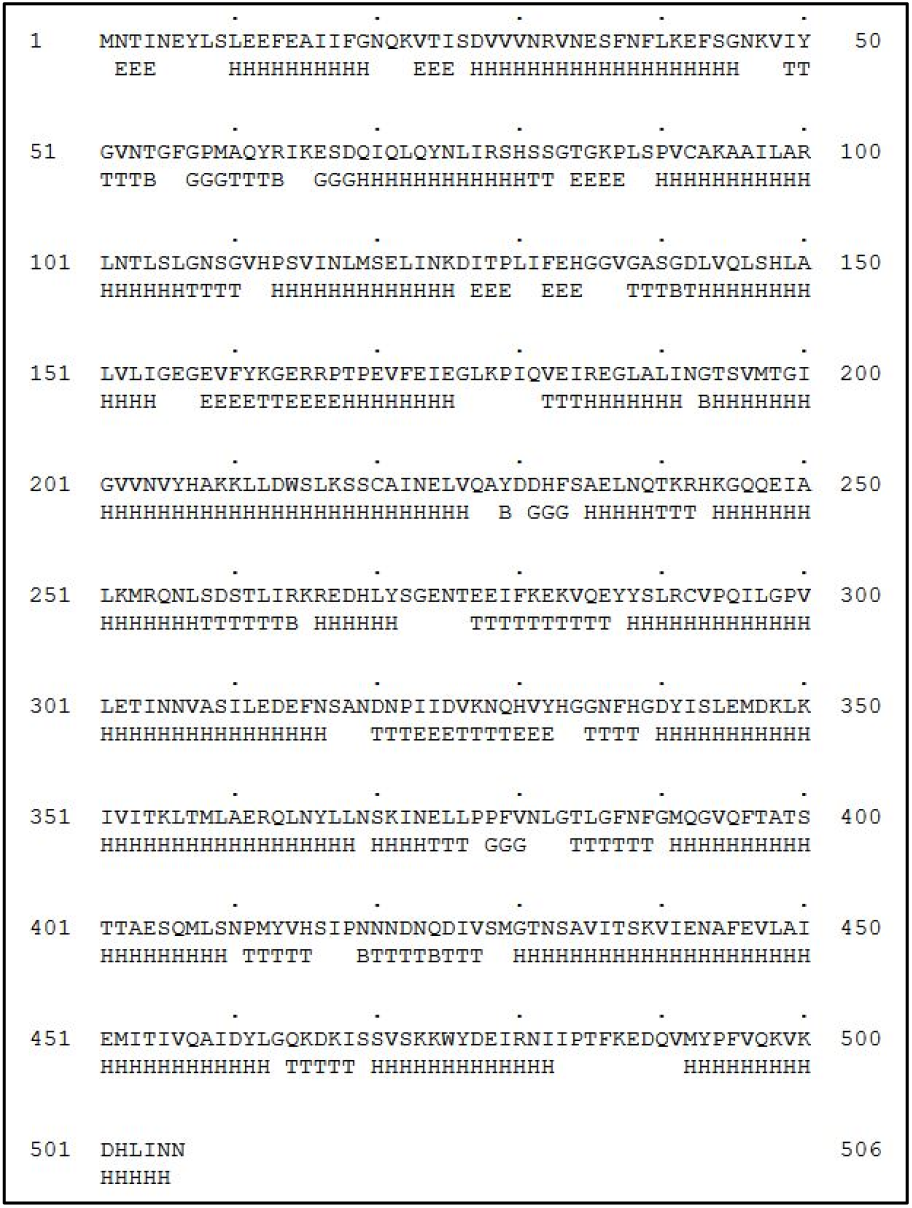
The secondary structures retrieved from FjTAL_1_AF2 as one of the top candidate structures. **H**: Alpha helix, **G**: 3-10 helix, **I**: PI-helix, **E**: Extended conformation (beta strand), **B**: Isolated bridge, **T**: Turn, **C**: Coil (none of the above)

**Fig.4.**
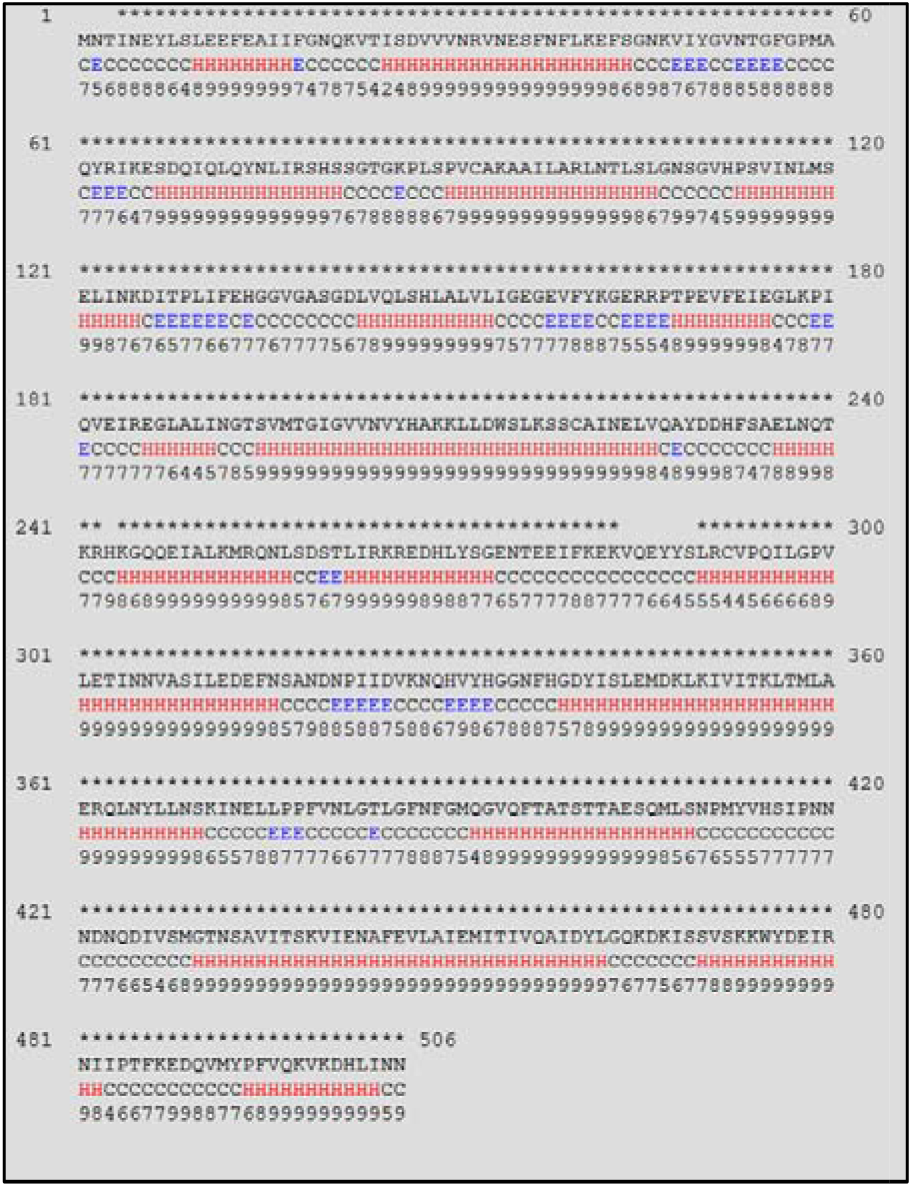
Secondary structures predicted by PROTEUS2. (C: coil, E: beta strand, H: alpha helix)

The structure of FjTAL_1_AF2 is shown in Fig.5 as an example.^25,26^

**Fig.5.**
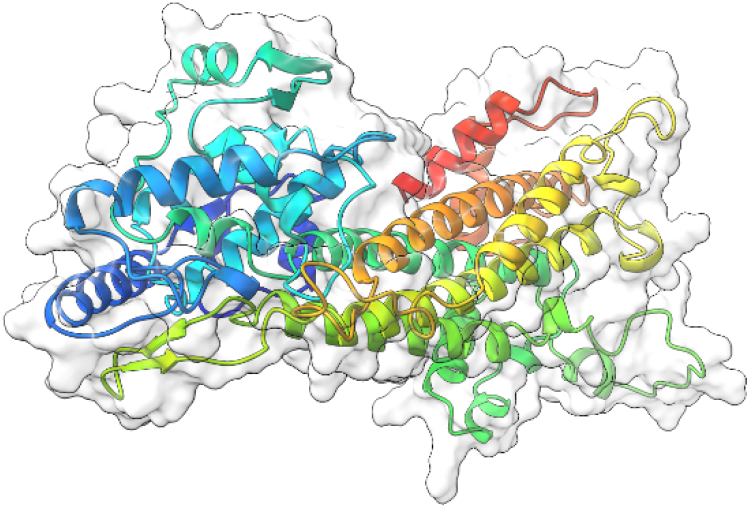
Ribbon visualization of FjTAL_1_AF2.pdb as one of the candidate structures for FjTAL.

We also passed the pdb files to BIOVIA Discovery Studio in order to calculate RMSD, as a measure of similarity among the structures (see Table4 and Fig.6).

**Fig.6.**
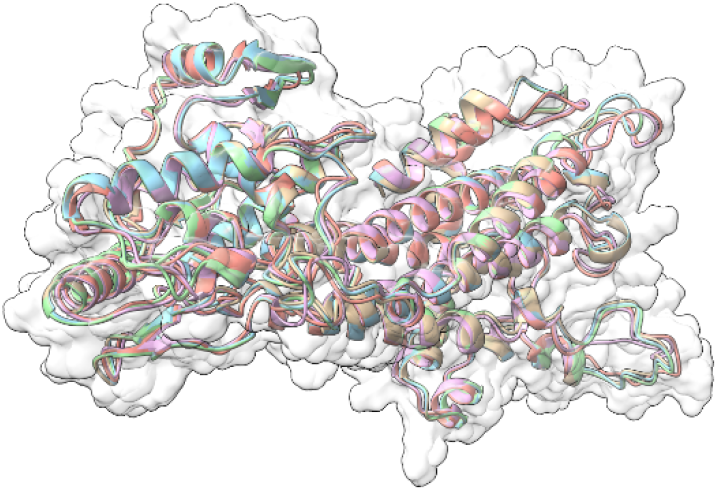
The superimposed structure of FjTAL

**Table4.** RMSD values for each pair of models. We have arbitrarily taken FjTAL_1_AF2 as the reference model.

| Molecule | C-Alpha | Main-chain | Side-chain | All Protein |
| --- | --- | --- | --- | --- |
| FjTAL_3_AF2 | 0.237 | 0.285 | 0.67 | 0.512 |
| FjTAL_3_Rob | 1.456 | 1.48 | 2.608 | 2.112 |
| FjTAL_4_AF2 | 0.276 | 0.313 | 0.736 | 0.563 |
| FjTAL_4_Rob | 1.423 | 1.447 | 2.608 | 2.1 |

Note that there exists a significant difference (ΔRMSD > 2 Angstrom) in the structures predicted by Robetta/RoseTTAFold and ColabFold/AlphaFold2. Their difference mainly stems from the coils and turns rather than the alpha helices. Without the availability of an experimentally-verified 3D structure for the enzyme, such workflow was the only logical way one could turn into to find out more about the structure of FjTAL and its relevance to its function.

### 3.3 FjTAL Molecular Docking to L-Tyrosine, p-Coumaric Acid and Caffeic Acid

With the availability of the predicted structure of FjTAL, a molecular docking task was executed. The structures of ligands i.e., L-tyrosine, p-coumaric acid and caffeic acid were retrieved from ChemSpider and their structures were subsequently adjusted^27–30^ to pH=7.4. Then, an MM2 minimization task was performed to retrieve the best energetically-favored representation of the ligands. The five receptor pdb files were docked with three potential ligands, L-tyrosine, p-coumaric acid and caffeic acid via the vina^31^-based docking server, CB-Dock^32,33^.

CB-Dock reports five cavities and their corresponding vina score for each pair of ligand and receptor. The top results, all of which possessing a vina score =< −5.5, were selected for further investigations.

We used mol*^34^ to determine the residues that constituted the binding pocket of FjTAL in each of these complexes. Interestingly two distinct binding sites, named after their first residues as VQAY and VIY cavities, consistently appeared in most of the structures (see Table 5, Fig.7 and Fig.8).

**Fig.7.**
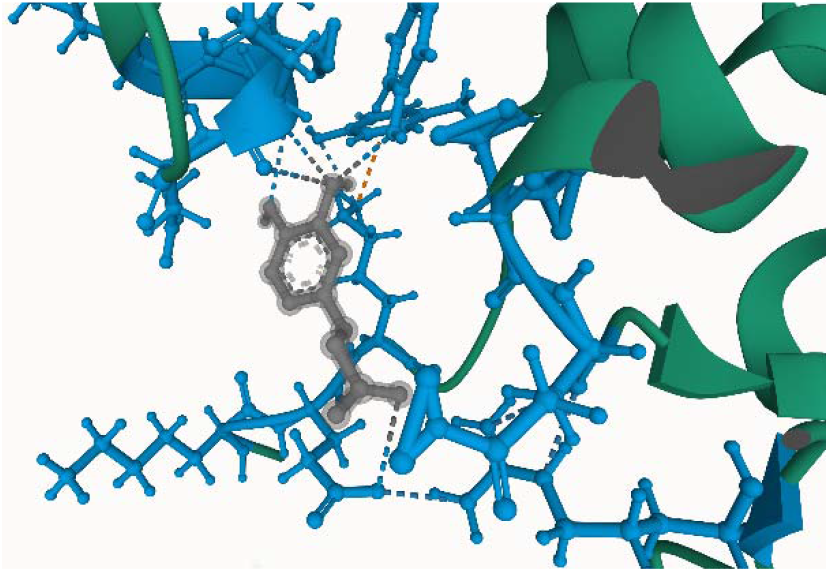
The predicted binding site of pH-adjusted, MM2 minimized, caffeic acid and FJTAL_1_AF2.pdb with vina score = −5.8 The blue residues depict the VQAY pocket, which is comprised of less evolutionary conserved residues.

**Fig.8.**
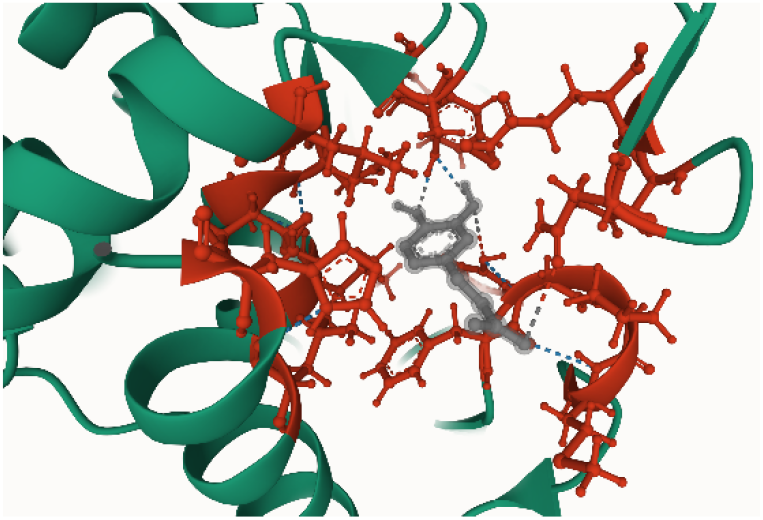
The predicted binding site of pH-adjusted, MM2 minimized, caffeic acid and FjTAL_1_AF2.pdb with vina score = −5.9 The red residues depict the VIY pocket, consisted of many evolutionary conserved residues.

**Table 5.** The molecular docking top results and their corresponding binding cavities. VQAY cavity is in blue and VIY cavity is in red. Other colors represent rare and presumably false binding sites.

|  | L-tyrosine |  | p-coumaric acid |  | caffeic acid |  |
| --- | --- | --- | --- | --- | --- | --- |
| FjTAL_1_AF2 | -6.2 | -5.7 | -5.7 | -5.6 | -5.9 | -5.8 |
| FjTAL_3_AF2 | -6.0 | -5.7 | -5.6 | -5.6 | -5.8 | -5.8 |
| FjTAL_3_Rob | -5.7 |  | -6.0 |  | -6.1 | -5.6 |
| FjTAL_4_AF2 | -6.1 | -5.7 | -5.7 | -5.6 | -5.9 | -5.6 |
| FjTAL_4_Rob | -5.5 |  | -5.5 |  | -5.7 |  |

Some of the residues that constitute the VIY cavity, are known to form the prosthetic group, 3,5-dihydro-5-methylidene-4H-imidazol-4-one (MIO), which has been proved to act as an electrophilic species in the deamination reaction catalyzed by Phenylalanine Ammonia Lyase (PAL).^35^ VIY cavity homologs also appeared in the experimentally-verified structure of *Rhodobacter sphaeroides* Tyrosine Ammonia Lyase, RsTAL in complex with caffeate and coumarate^36^. The VQAY cavity exhibits very few evolutionary-conserved residues, suggesting that it is either completely an artifact or it could be another binding site to the ligands as well as the VIY cavity. Site directed mutagenesis studies can help us further illustrate the importance of VQAY cavity.

The molecular docking results demonstrate that pHCA and caffeic acid bind to FjTAL in the same pocket as L-tyrosine, supporting the idea that these two may competitively inhibit the activity of FjTAL, and further result in a negative feedback response.^6^

## 4. Conclusion

In this study, the 3D structure of *Flavobacterium johnsoniae* Tyrosine Ammonia Lyase (FjTAL) has been predicted by employing the state of art computational tools. The candidate structures were filtered and subsequently docked with three potential ligands, L-tyrosine, p-coumaric acid and caffeic acid. The results indicated that two potential binding cavities may exist on the surface of FjTAL, one of which exhibited great sequence identity with the experimentally-verified docking site of RsTAL. The other detected binding site may be a false positive result or a second binding site for the ligands. Eventually, site directed mutagenesis studies and X-ray crystallography data would be extremely beneficial to help us understand the structure of FjTAL comprehensively.

## Supplementary material

The PDB files of the predicted structures, ligands and docking results are uploaded.

## Notes

### Competing Interest Statement

The authors have declared no competing interest.

### Summary of Updates

Name has been edited. A few sentences were added to some of the parts of the manuscript.

